# Dual role for fungal-specific outer kinetochore proteins during cell cycle and development in *Magnaporthe oryzae*

**DOI:** 10.1101/400549

**Authors:** Hiral Shah, Kanika Rawat, Harsh Ashar, Rajesh Patkar, Johannes Manjrekar

**Affiliations:** Bharat Chattoo Genome Research Centre, Department of Microbiology and Biotechnology Centre, The Maharaja Sayajirao University of Baroda, Vadodara 390002, Gujarat, India; Biotechnology Programme, Department of Microbiology and Biotechnology Centre, The Maharaja Sayajirao University of Baroda, Vadodara 390002, Gujarat, India; Centre for Ecological Sciences, Indian Institute of Science, Bengaluru-560012, Karnataka, India

**Keywords:** Kinetochore, Dam1 complex, Filamentous fungus, *Magnaporthe*, Rice Blast

## Abstract

The outer kinetochore DASH/DAM complex ensures proper spindle structure and chromosome segregation. While DASH complex protein requirement diverges among different yeasts, its role in filamentous fungi has not been investigated so far. We studied the dynamics and role of middle (Mis12) and outer (Dam1 and Ask1) kinetochore proteins in the filamentous fungal pathogen, *Magnaporthe oryzae,* which undergoes multiple cell cycle linked developmental transitions. Both Dam1 and Ask1, unlike Mis12, were recruited to the nucleus specifically during mitosis. While Dam1 was not required for viability, loss of its function (*dam1*Δ mutant) delayed mitotic progression, resulting in impaired conidial and hyphal development in *Magnaporthe*. Intriguingly, both Dam1 and Ask1 also localised to the hyphal tips, in the form of punctae oscillating back and forth from the growing ends, suggesting that *Magnaporthe* DASH complex proteins may play a non-canonical role in polarised growth during interphase, in addition to their function in nuclear segregation during mitosis. Impaired appressorial (infection structure) development and function in the *dam1*Δ mutant suggest that fungus-specific Dam1 complex proteins could be an attractive target for a novel anti-fungal strategy.

**Summary Statement:** DASH complex proteins are differentially recruited to the nucleus during cell division and are intriguingly involved in polarised growth during development and differentiation in the rice blast fungus.

## Introduction

Chromosome segregation is a determining step in cell division. During mitosis, the kinetochore (KT) brings about precise chromosome segregation at anaphase and anomalies in chromosome separation are associated with malfunction and disease in multicellular organisms. In unicellular yeasts, such aberrant chromosomal segregation often leads to loss in viability. The KT varies in structure and composition among eukaryotes (Van Hooff et al., 2017). The microtubule (MT)-associated DAM/DASH Complex is a fungus-specific component of the outer kinetochore. This complex varies in its role in the yeasts *Saccharomyces cerevisiae, Schizosaccharomyes pombe* and *Candida albicans*, where it has been studied extensively; however only very little is known about its function in filamentous fungi.

The DASH complex is made up of 10 subunits, namely <u>D</u>am1, <u>A</u>sk1, <u>S</u>pc34, <u>H</u>sk3, Dad1-4, Duo1 and Spc19, all of which are essential for viability in *S*. *cerevisiae* and *C*. *albicans*. On the contrary, none of the members is individually essential for survival in *S*. *pombe*. In *S*. *cerevisiae,* Dam1 is required for proper spindle assembly and elongation (Hofmann et al., 1998, Cheeseman et al., 2001), while Ask1 is required for bipolar attachment (Janke et al., 2002). Dam1 directly interacts with MTs and other DASH members to form DASH complex oligomers (Legal et al., 2016). The yeast DAM complex members are MT plus-end interacting proteins (TIPs). The *S*. *pombe dam1*Δ mutant displays chromosome segregation delay and ectopic septation (Sanchez-Perez et al., 2005). Further, *S*. *pombe* DASH complex mutants are sensitive to the MT poison thiabendazole, low temperature and high osmolarity - all phenotypes shared by genes involved in MT stability and dynamics (Sanchez-Perez et al., 2005, Gao et al., 2010). Dam1 was also discovered as a multicopy suppressor of mutations in the spindle regulator Cdc13, and MT plus-end binding protein Mal3/Eb1 (Sanchez-Perez et al., 2005). Further, in *S*. *pombe*, retrieval of unclustered kinetochores is dependent on Dam1 (Franco et al., 2007).

In *S*. *pombe*, Dam1 is recruited to the KT in early mitosis, where it is observed as a single spot that segregates into several spots at metaphase, while Dad1 remains associated with the KT throughout the cell cycle. The DASH complex spots co-localise with other KT proteins like Mtw1 (Mis12) and Ndc80 (Sanchez-Perez et al., 2005). In *S*. *cerevisiae*, Dam1 is present at the KT throughout the cell cycle, clustered as a single nucleus-associated spot during interphase, which divides into two spots closely associated with the spindle pole bodies at mitosis. Dam1 has also been observed all along the spindle in *S*. *cerevisiae*. Association of Ask1 with the KT is dependent on Ndc10 and MT spindle. *In vitro* microtubule binding studies using *S*. *cerevisiae* proteins have shown 16-member oligomeric rings that encircle the MT. However, rings are not essential for MT attachment and evidence for such rings *in vivo* is lacking so far.

With respect to nuclear envelope continuity, fungi exhibit a whole spectrum of mitoses. While *S*. *cerevisiae* and *S*. *pombe* both undergo closed mitosis, filamentous fungi show differences; semi-open in *Aspergillus nidulans*, open in *Ustilago maydis* and semi-closed in *Magnaporthe oryzae*. Further, in *S*. *cerevisiae* mitosis lasts around 50 minutes while in filamentous fungi it is completed in less than 5 minutes. In addition, mitosis in filamentous fungi is often associated with distinct morphological differentiation not observed in most yeasts, setting different requirements and constraints on spindle assembly and elongation. Thus, owing to these differences, it is likely that KT-spindle structure and assembly differ in filamentous fungi. Unlike yeasts, the filamentous fungal pathogen *M*. *oryzae* undergoes several morphologically distinct developmental transitions during its life cycle.

During the *M*. *oryzae* pathogenic life cycle, the three-celled conidium germinates on the leaf surface to extend a polarised germ tube. While entry of the germinating cell nucleus into S phase is required for the swelling of the germ tube tip (switching from polarised to isotropic growth to form the incipient appressorium), entry into mitosis is necessary for the further development of the appressorium (Saunders et al., 2010a). The diploid nucleus of the germinating cell undergoes the first round of infection-related mitosis in the germ tube. Nuclear division is followed by migration, during which one daughter nucleus returns to the germinated cell in the conidium and the other travels into the appressorium. Subsequently, exit from mitosis is required for appressorium maturation and function (Saunders et al., 2010b). Interestingly, the appressorium then switches back to polarised growth to form the penetration peg that breaches the host epidermis and elaborates into the primary infection hypha (IH). This developmental transition is again dependent on S phase checkpoints. At this point, the second round of infection-related mitosis in the mature appressorium contributes a nucleus to the IH. The IH nuclei undergo semi-closed mitosis lasting roughly three minutes (Jones et al., 2016). Thus, the crucial morphological transitions during *M*. *oryzae* infection are tightly coupled to different stages of the cell cycle. Secondary hyphae then develop from the primary IH and spread to the neighbouring host cells, giving rise to typical disease lesions in a few days. These secondary hyphae later also give rise to the aerial hyphae, some of which eventually form conidiophores bearing 3-5 sympodial conidia. A single lesion can produce a new generation of thousands of conidia that can initiate a new infection cycle.

*M*. *oryzae* infects rice and several other cereal crops across the world, and is a serious threat to global food security (Dean et al., 2012). Investigating the role of fungus-specific components in the development of this fungus would provide new targets for novel anti-fungal strategies. DASH complex proteins are not found in rice or other plant hosts making them potential candidates; however, within filamentous fungi, only the Duo1 subunit of *Magnaporthe* has been studied so far. The *M*. *oryzae* Duo1 plays a role in conidiation and full virulence in rice, but not much information is available regarding its function as a kinetochore protein in mitosis or its localisation during the cell cycle (Peng et al., 2011). In this study, we focussed on the role of DASH complex proteins in chromosome segregation, especially during the cell cycle-regulated morphological transitions in *M*. *oryzae*. Using deletion mutants and GFP-tagged strains, we studied the behaviour of Dam1 and Ask1 during nuclear division, spindle dynamics and nuclear migration, and its effect on *M*. *oryzae* development. We show that in addition to its role in proper chromosome segregation, Dam1 - through its localisation in the hyphal tip compartment during interphase – is required for normal conidiation and polarised hyphal growth.

## Results

### DASH complex protein Dam1 is recruited to the nucleus during mitosis

We identified the orthologue of *S*. *cerevisiae* and *S*. *pombe* Dam1 in *M*. *oryzae* (MGG_00874) using BLASTP. Although the overall protein sequence similarity was ~30-40%, the DASH complex domain of *M*. *oryzae* Dam1 showed significant homology (~60%) with that of yeasts. The Dam1 proteins show considerable size variation, ranging from 343 amino acids in *S*. *cerevisiae* to 121 in *Cryptococcus neoformans*, with the *M*. *oryzae* protein being of 220 amino acids (Table S1). The protein size variation was due to differences in the length of the C-terminal region. We further identified another member of the DASH complex, Ask1 (MGG_07143), which is known to directly interact with Dam1 in *S*. *cerevisiae*. To study the subcellular localisation and dynamics with respect to that of the nucleus, we tagged Dam1 and Ask1, separately, with GFP (green fluorescent protein) in a strain expressing histone H1-mCherry (hH1-mCherry). As a marker for kinetochore position, we also studied the localisation of the middle kinetochore MIND Complex protein Mis12 (MGG_06304) in *Magnaporthe*.

In the vegetative hyphae, Mis12-GFP was observed as a single distinct puncta at the periphery of the interphase nucleus (arrowhead; Fig. 1A). In contrast, the DASH complex proteins GFP-Dam1 and Ask1-GFP did not localise predominantly to the nucleus during interphase (Fig. 1A). Occasionally, GFP-Dam1 or Ask1-GFP punctae were seen in the cytoplasm along the hyphae (arrows; Fig. 1A). We presumed these were DASH complex proteins associated with the cytoplasmic microtubules. At the onset of mitosis (marked by chromosome condensation), while the Mis12-GFP de-clustered, the GFP-Dam1 punctae appeared at the nucleus and persisted there throughout the mitotic process and the subsequent nuclear migration (arrowheads, Fig. 1B). At the end of mitosis, the Mis12-GFP and GFP-Dam1 spots re-clustered into a single spot per nucleus (Fig. 1B). To determine whether DASH complex proteins had similar dynamics during the early infection process of *M*. *oryzae*, we studied GFP-Dam1 or Ask1-GFP localisation during appressorium formation. The Ask1-GFP and GFP-Dam1 punctae were associated with the nucleus during nuclear division and migration in the course of appressorial development (lower panels; Fig. 1B and S2). Thus, while the middle KT layer protein Mis12 was constitutively associated with the *M*. *oryzae* kinetochore, the outer layer DASH complex proteins were specifically recruited to the nucleus during mitosis.

**Figure 1:**
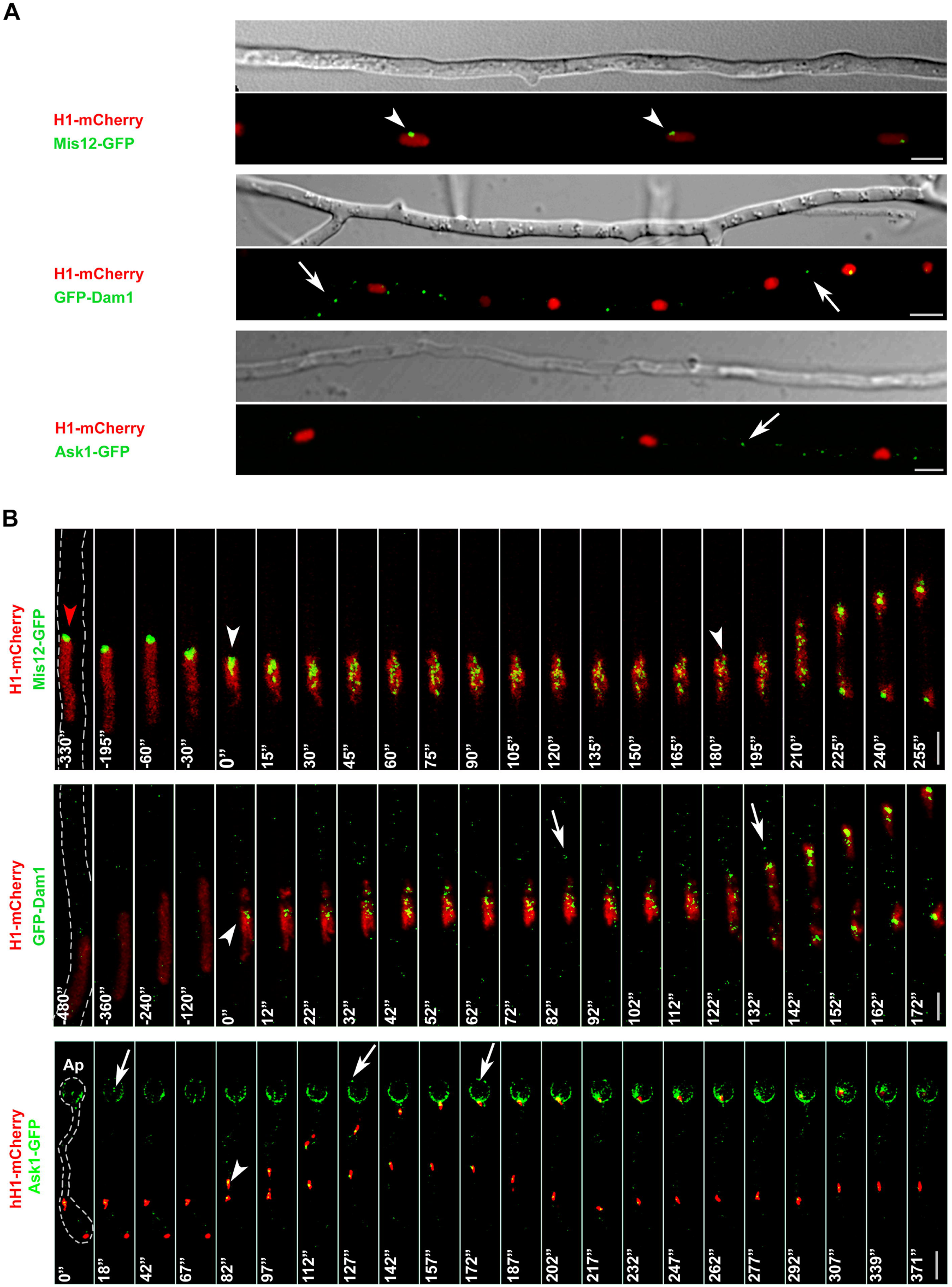
Dam1 is recruited to the nucleus at the onset of mitosis in *M*. *oryzae*. A) Localisation of middle and outer KT proteins, Mis12-GFP, GFP-Dam1 and Ask1-GFP, during interphase in vegetative hyphae (Bars = 5 µm). B) Time-lapse images showing dynamics of Mis12-GFP or GFP-Dam1 during mitosis in vegetative hyphae (Bars = 3 µm) or Ask1-GFP during appressorium (indicated as Ap) development (Bar = 10 µm). Fungal structures are marked with dashed outlines. Arrowheads indicate nuclear-associated Mis12-GFP or GFP-Dam1 or Ask1-GFP. Red arrowhead indicates Mis12-GFP associated with the nucleus prior to mitosis. Arrows denote GFP-Dam1 or Ask1-GFP spots likely associated with cytoplasmic microtubules along the vegetative hyphae or in the appressorium.

### Dam1 plays a key role in proper segregation of chromosomes

Dynamics of the nuclei marked with hH1-mCherry and MTs or spindle (β-Tubulin-sGFP) were monitored during mitosis in the vegetative hyphae of *M*. *oryzae*. Alongside, to investigate the role of DASH complex proteins in mitosis, we generated *DAM1* deletion strains, hereafter referred to as *dam1*Δ, in the WT or the strain expressing hH1-mCherry and Tub-GFP. We deleted *ASK1* in the GFP-Dam1 expressing strain (hereafter referred to as *ask1*Δ). At the onset of mitosis in the WT strain, prophase was marked by microtubule re-organisation and chromosome condensation. It was characterised by loss of cytoplasmic microtubule arrays and establishment of a single tubulin focal point at the nucleus, likely a site for spindle microtubule nucleation. This was followed by the assembly of an intensely fluorescent bipolar spindle, which underwent re-orientation to align along the long axis of the hypha. Once formed, the spindle length remained largely constant until spindle elongation during anaphase. The chromosomes then segregated into two daughter nuclei that migrated to opposite ends (Fig. 2A). In the absence of Dam1 function, the *dam1*Δ strain showed longer spindles (6.47 ± 0.26 μm) when compared to the WT (4.12 ± 0.19 μm) (Fig. 2B). We further determined the time for which the spindle persisted until the onset of anaphase in the WT and *dam1*Δ strains. The *dam1*Δ mutant showed a considerable delay in spindle elongation as compared to the WT (Fig. 2C). While chromosome segregation was mostly initiated within 3 minutes in the WT (2.13 ± 0.34 mins), anaphase onset in several *dam1*Δ mutant cells took more than 10 minutes (12.87 ± 2.27 mins) (Fig. 2D). The time spent in metaphase or prior to anaphase varied considerably between individual cells in the *dam1*Δ strain (3-39 minutes) (Fig. 2D). The duration of mitosis in the germ tube during appressorium development showed a similar delay in the *dam1*Δ mutant. Importantly, in a few cells, we occasionally observed unequal segregation of nuclear material or lagging chromosomes that moved behind the rest (arrows, Fig. 2C). Thus, Dam1 function likely ensures correct spindle structure and separation of chromosomes at anaphase, allowing proper mitotic progression during both hyphal growth and appressorial development in *M*. *oryzae*.

**Figure 2:**
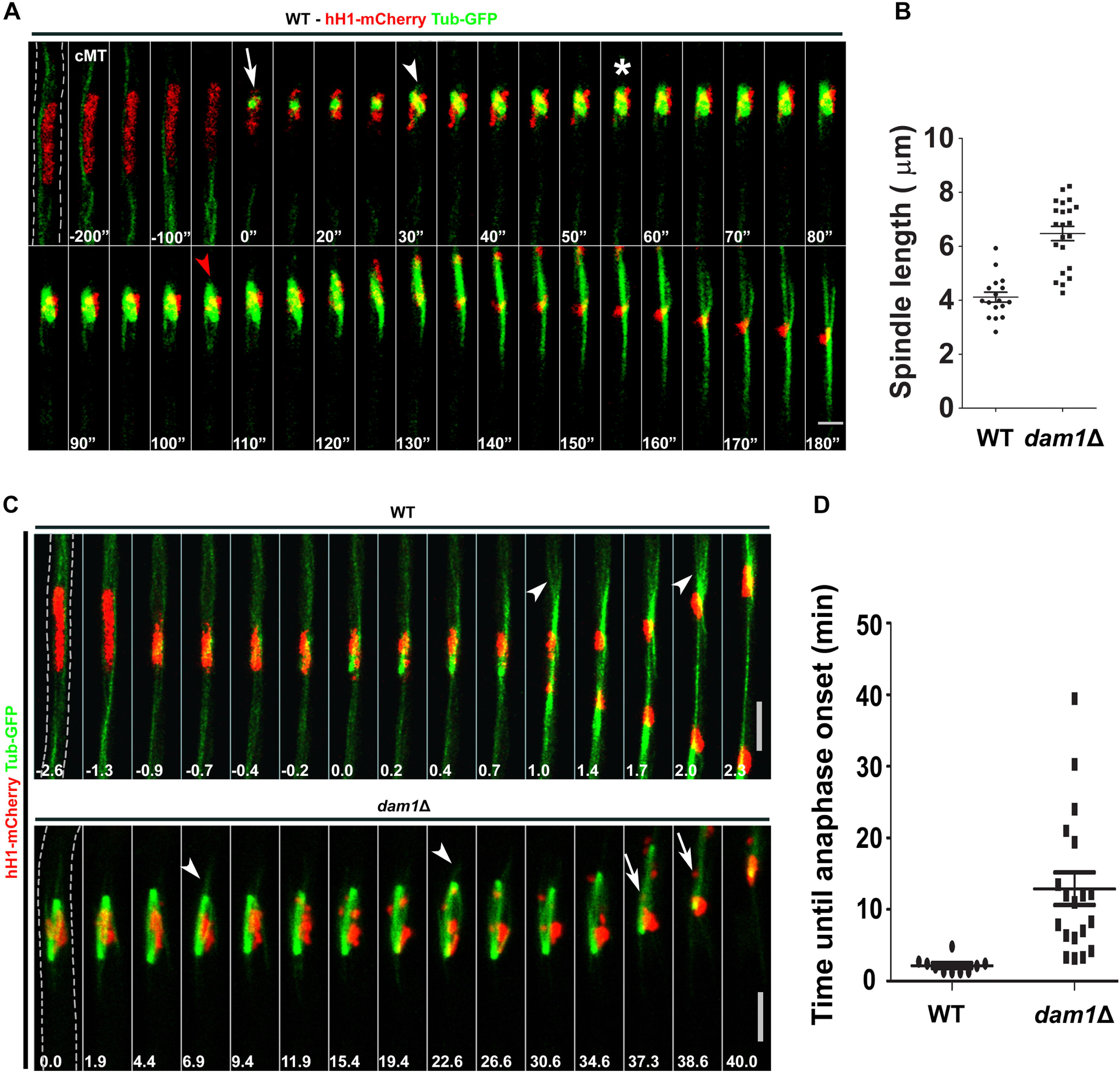
Dam1 plays an important role in proper segregation of chromosomes during nuclear division. A) Dynamics of the mitotic spindle in the WT *M*. *oryzae* marked with Tub-GFP and hH1-mCherry. Arrow depicts entry into mitosis, arrowhead indicates assembly of a bipolar spindle, asterisk shows spindle re-orientation, and red arrowhead indicates onset of spindle elongation, cMT-cytoplasmic MTs. Bars = 3 µm. Hyphal borders are indicated with dashed lines. B) Scatter Plot showing spindle length in the WT or *dam1*Δ *M*. *oryzae* during mitosis. Data represents mean + S.E.M. with n_WT_ = 17, n_*dam1*Δ_ = 22, *P*< 0.05. C) Time-lapse images of mitosis in the WT or *dam1*Δ vegetative hyphae. Bars = 5 µm. Numbers indicate time in minutes. Hyphae are marked with dashed outlines. Arrowheads indicate astral MTs and arrows show lagging chromosomes. D) Scatter Plot showing time until spindle elongation in anaphase in WT or *dam1*Δ during mitosis in vegetative hyphae. Data represents mean + S.E.M. with n_WT_ = 11, n_*dam1*Δ_ = 19, *P*< 0.05.

### The DASH complex is involved in polarised vegetative hyphal growth

Next, we looked at the physiological effects of delayed mitosis due to the loss of Dam1 function. We found that the *dam1*Δ (Fig. 3A) and *ask1*Δ (Fig. S1) strains had a significantly reduced (upto 60%) colony diameter mainly due to aberrant vegetative hyphal extension and morphology, suggesting impaired cell polarity in the mutant strains of *M*. *oryzae* (Fig. 3B). In *M*. *oryzae*, hyphae grow in a monopolar fashion during apical extension and lateral branching. The branches arise close to septa at acute angles to the growing primary hypha at fairly regular intervals. The hyphae of the *dam1*Δ strain showed more frequent branching and a zigzag or curved morphology, unlike the mostly straight hyphae of the WT (Fig. 3B). Further, calcofluor white (CFW) staining of the *dam1*Δ hyphae showed irregularly sized cell compartments compared to the uniform compartments in the WT. The *dam1*Δ mutants showed some backward branches (opposite to the direction of primary hyphal growth) and tip splitting – both phenotypes that are rarely seen in the WT.

**Figure 3:**
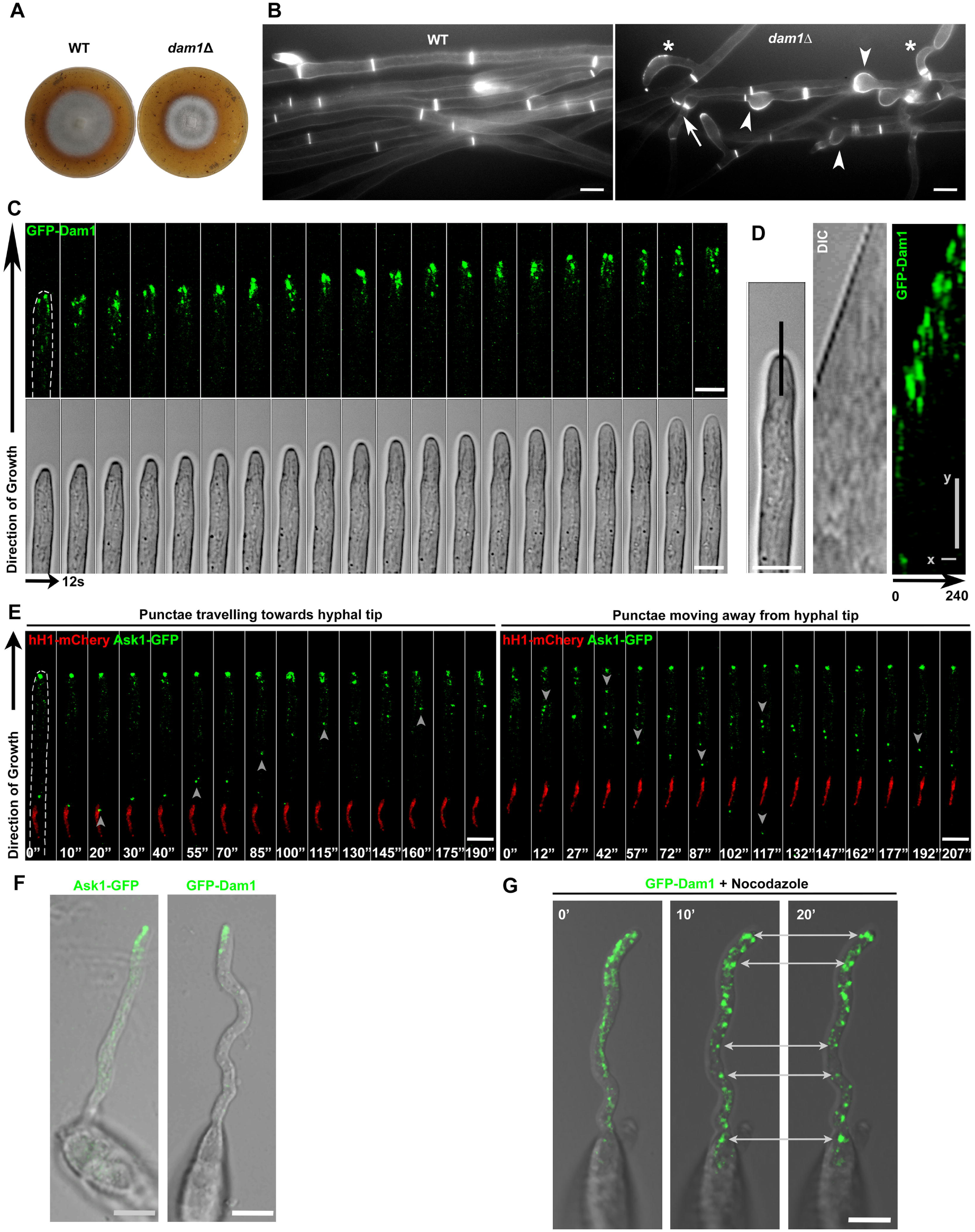
Dam1/DASH Complex is involved in polarised vegetative hyphal growth during interphase. A) WT or *dam1*Δ vegetative hyphae stained with calcofluor white (CFW). Arrowheads, arrows and asterisks indicate bulged hyphae, multiple branching and curved hyphae, respectively, in the *dam1*Δ, unlike the regular hyphae seen in the WT. Bars = 5 µm. B) Vegetative growth of the WT or *dam1*Δ strain at 8 dpi on prune agar. C) Localisation of GFP-Dam1 at the hyphal tip during vegetative growth. Hyphal borders are shown with dashed lines. Bars = 5 µm. D) Kymographs of time-lapse images of GFP-Dam1 shown in (C). Kymographs are plotted along the black line marked on the hyphal tip for 240 seconds. Scale Bars: x = 48s, y = 1.5 µm. E) Oscillation of Ask1-GFP from vegetative hyphal tip during interphase. Arrowheads mark the Ask1-GFP punctae moving towards and away from the tip. F) Ask1-GFP or GFP-Dam1 localise to the germ tube tip during polarised growth on a hydrophobic surface. Effect of nocodazole treatment on GFP-Dam1 localisation is depicted. Arrows mark Ask1-GFP or GFP-Dam1 at the germ tube tip. Bars = 5 µm.

Interestingly, we observed that GFP-Dam1 localises to the hyphal tip in the form of distinct spots during interphase (Fig. 3C). These intense spots were highly dynamic, and moved forward with the growing tips (Fig. 3D). To check whether this non-conventional localisation during interphase was specific to Dam1 or involved the DASH complex, we also studied the Ask1-GFP localisation. Indeed, Ask1-GFP too localised to the growing hyphal tips in a highly similar manner. Further, Ask1-GFP (Fig. 3E) and GFP-Dam1 spots showed an oscillatory movement to the tip and back towards the nucleus. Another stage of the *M*. *oryzae* life cycle that involves polarised growth is the extension of the germ tube during early stage of pathogenic development. Dynamic Ask1-GFP and GFP-Dam1 spots, similar to the ones seen in the hyphal tip, were observed at the tips of germ tubes, suggesting a role for the DASH complex during polarised growth in *M*. *oryzae* (Fig. 3F). To determine whether such localisation to the tips required an intact DASH complex (or at least that Dam1 did not behave completely independently of the rest of the DASH complex), we deleted *ASK1* in the GFP-Dam1 strain. We found that the intense punctae at the tips were lost in the absence of Ask1 function. Upon treatment with the microtubule-destabilising compound nocodazole, the dynamic GFP-Dam1 punctae became static aggregates, randomly distributed along the germ tube cytoplasm (Fig. 3G). Our observations show that the growth defects seen in the *dam1*Δ strain are a combined outcome of loss of Dam1 function during mitosis as well as polarised growth during interphase. Further, MT-based oscillatory behaviour of Dam1 suggests a non-canonical function for the DASH complex protein prior to the onset of mitosis in *Magnaporthe*.

### Conidiogenesis is marked by three distinct rounds of mitosis in *M*. *oryzae*

Production of three-celled conidia is a critical developmental step in the life cycle of *M*. *oryzae*. However, unlike in the other stages of the life cycle, the mitotic events and cell cycle regulation involved in conidial development have not been studied in great detail so far. Here, we first monitored mitosis during conidium development in the strain expressing hH1-mCherry and Tub-GFP. It has been already shown that conidial development starts with the shift from polarised to isotropic growth, where the tip of the aerial hypha starts swelling to form an incipient conidium (conidiophore) (Deng et al., 2009). We found that the first nuclear division took place in the stalk and was followed by nuclear migration where one of the daughter nuclei travelled into the conidium cell (mitosis I; Fig. 4A) and the other remained in the stalk. This was followed by cytokinesis, which likely occurred at the neck of the incipient conidium, away from the site of mitosis in the stalk. This spatial uncoupling of cytokinesis from mitosis was similar to the events observed during appressorium development, and different from what was observed in vegetative hyphal growth. The single-celled conidium then grew and elongated into an oval shaped cell with the nucleus positioned close to the centre. This nucleus underwent division, with one of the daughter nuclei remaining positioned at the base (towards the stalk) of the conidium and the other moving to the opposite end. The process was accompanied by active re-organisation of the microtubule network and septation at the site of mitosis to form an intermediate two-celled conidium (mitosis II; Fig. 4A). Subsequently, the nucleus in the second cell of the developing conidium underwent the third round of division, followed by cytokinesis to form the middle and terminal cells of the mature 3-celled conidium (mitosis III; Fig 4B).

**Figure 4:**
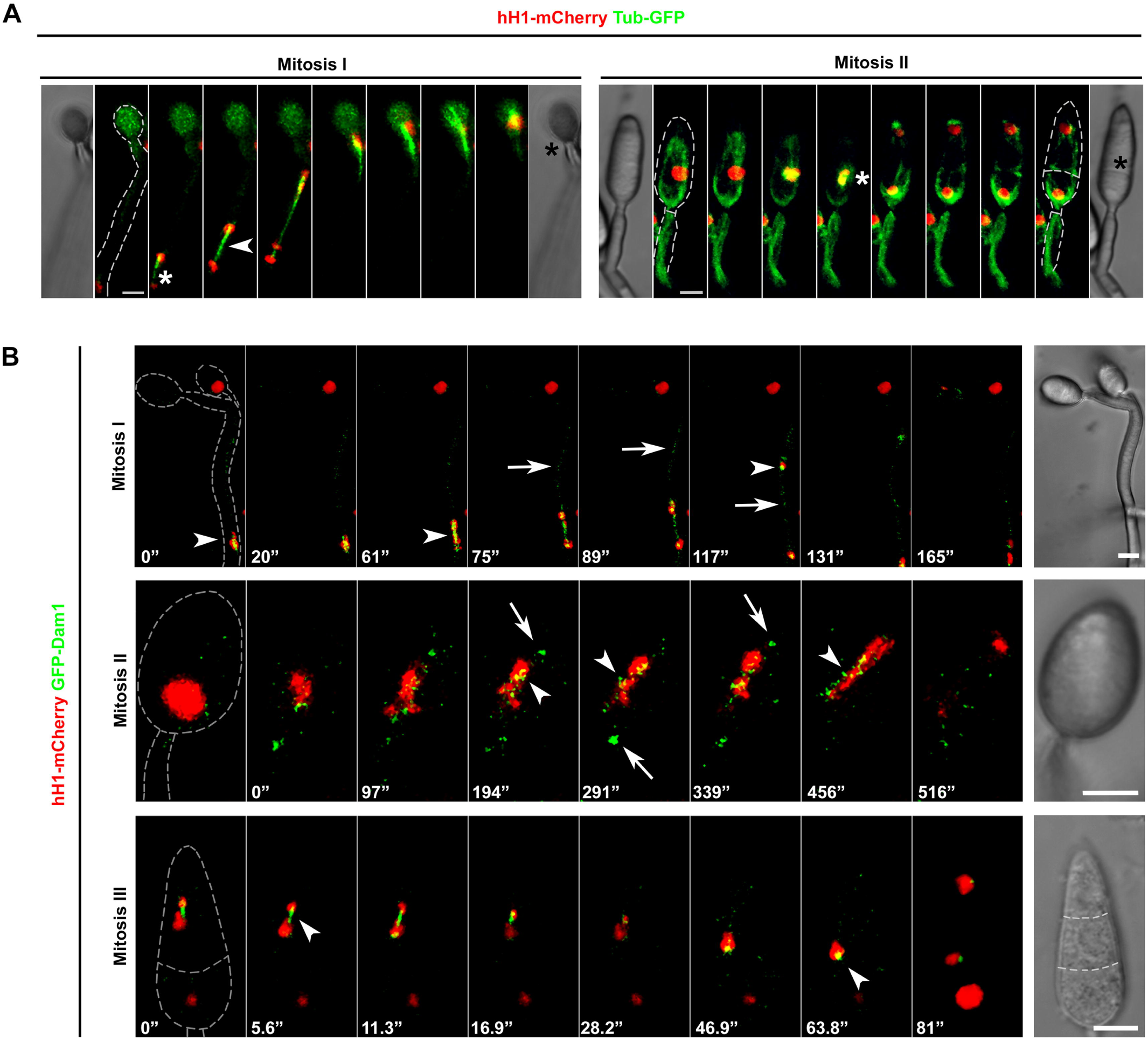
Conidium development is marked with three distinct rounds of mitosis. A) Dynamics of Tub-GFP-marked microtubules (MT) or spindle and hH1-mCherry-marked nuclear division during mitosis in the developing conidium. Mitosis I is observed in the stalk of the incipient conidium. Arrowheads mark the spindle. White asterisks indicate the site of mitosis while red asterisks denote the site of cytokinesis. B) Localisation of GFP-Dam1 during 3 successive rounds of mitosis in the developing conidium. Arrowheads depict nuclear-associated GFP-Dam1. Arrows indicate additional GFP-Dam1 spots. Fungal structures are shown with dashed grey outlines. Bars = 5 µm.

We next studied the localisation of GFP-Dam1 during mitoses in the developing conidia. We observed that GFP-Dam1 localised to the nucleus during all 3 rounds of nuclear division during conidiation (Fig. 4B). GFP-Dam1 appeared in the form of multiple spots during mitosis, and then clustered into two distinct spots, one per nucleus, and persisted till nuclear migration and positioning was complete. Dam1 or Ask1 did not associate with the nucleus during the intermittent interphases. Occasionally a cytoplasmic spot was seen during the initial conidiophore development prior to the first round of mitosis. Thus, Dam1 function plays a key role in three sequential and essential rounds of mitosis to form a complete three-celled asexual conidium.

### Dam1 function is crucial for proper conidiation

In addition to reduced vegetative growth, the *dam1*Δ strain showed flat and white colonies, in contrast to the fluffy, grey growth of the WT, likely due to defects in the development of aerial hyphae that give rise to conidia. To assess the role of Dam1 protein during asexual conidium development, we studied the growth and morphology of the aerial hyphae and conidiophores in the *dam1*Δ strain. Most of the conidiophores of the *dam1*Δ strain bore only 1 to 3 conidia as compared to the sympodial cluster of 3 to 5 conidia observed in the WT (Fig. 5A and B) after 24h of photo-induction. The total number of conidia in the *dam1*Δ strain was reduced to ~10% of the WT, and the mutant conidia were smaller (14 μm) in length than those in the WT (22 μm). In addition, the *dam1*Δ mutant showed aberrant morphology compared to the WT when stained with CFW. In contrast to the three-celled pyriform conidia seen in the WT, *dam1*Δ mostly produced one- or two-celled oval conidia (Fig. 5C and D). Similarly, the *ask1*Δ mutant too displayed defects in the total conidium number and morphology (Fig. S1). The WT three-celled conidia showed distinct cell boundaries with one nucleus per cell and an intense microtubular network, especially along the septa, likely indicative of septal microtubule organising centres (MTOCs). We further found that ~50% of the *dam1*Δ conidia had aberrant or collapsed nuclear and microtubular structures when compared to the WT (Fig. 5E and F). We infer that Dam1 function, likely by ensuring precise and orderly mitotic progression, is required for proper conidial development and morphology in *M*. *oryzae*.

**Figure 5:**
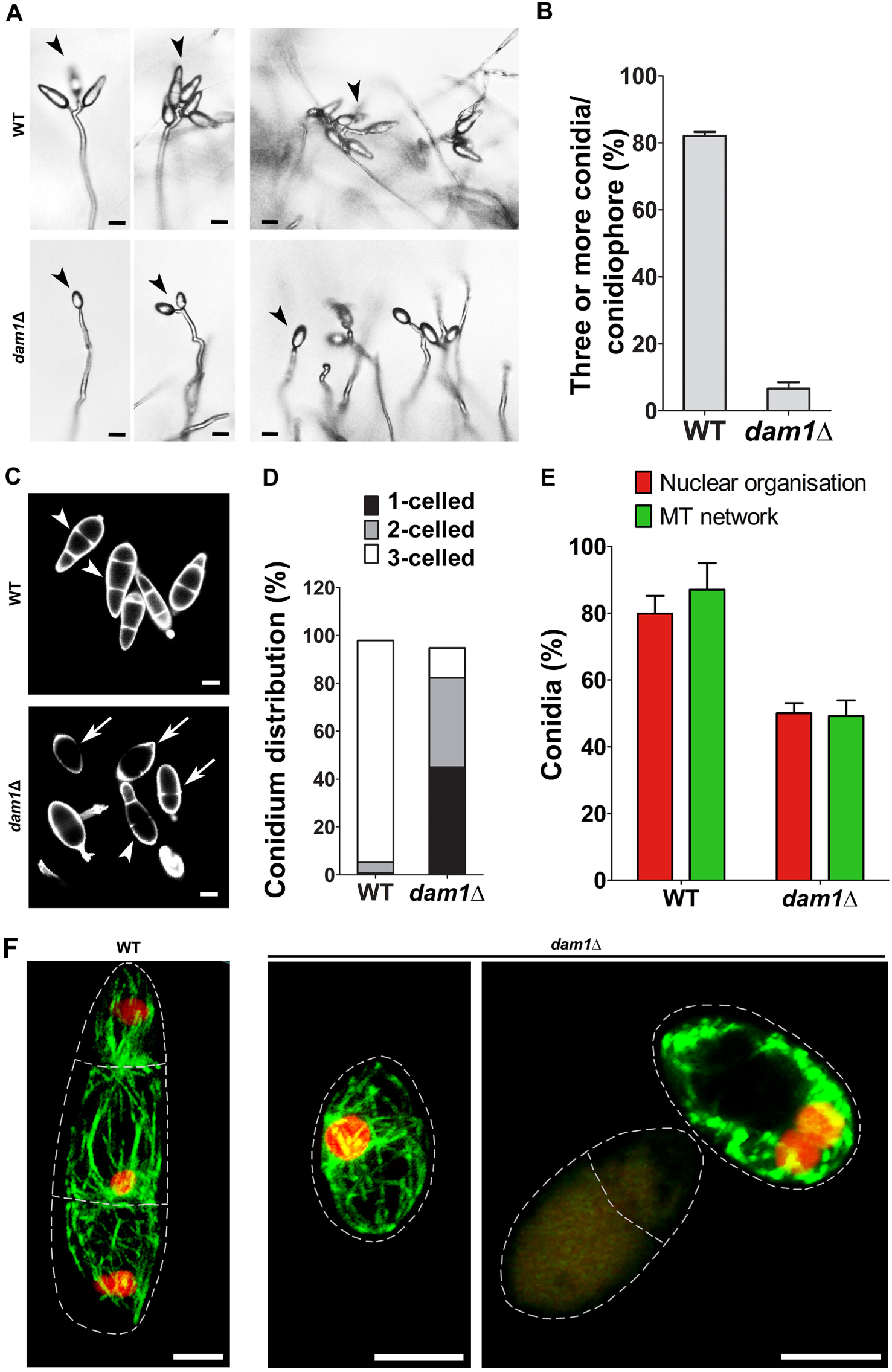
Loss of Dam1 function significantly alters conidiation. A) Difference in the conidiophore morphology and number of sympodial conidia (arrows) between the *dam1*Δ mutant and WT. Bars = 10 µm. B) Frequency of conidiophores bearing 3 or more conidia in the *dam1*Δ compared to the WT. *P*< 0.0001, t test, n = 300. C) Morphology of the *dam1*Δ or WT conidia stained with CFW. Arrowheads depict 3-celled WT-like conidia and arrows indicate 1- or 2-celled conidia. D) Bar chart showing frequency of conidia with different cell numbers (1, 2 or 3) in the WT or *dam1*Δ strain. *P*< 0.0001, t test, n = 300. E) Quantification of conidia with intact/normal nuclear and MT organisation in the WT or *dam1*Δ mutant, *P*< 0.05, t test, n = 100. F) Nuclear organisation and MT network in the WT or *dam1*Δ conidia expressing hH1-mCherry and Tub-GFP. Bars = 5 µm. Data represents mean + S.E.M. from three independent experiments.

### Dam1 function is required for proper pathogenic development and virulence

We studied the infection-related (appressorial) development in the *dam1*Δ strain, to assess the role of Dam1 in pathogenesis of *M*. *oryzae*. In *in vitro* assays, the majority of the *dam1*Δ conidia failed to germinate (Fig. 6A). While 81% of the WT conidia formed appressoria, only 20% of the *dam1*Δ mutant conidia formed the infection structure (Fig. 6A and B). A few mutant conidia developed aberrant germ tubes and/or appressoria, in contrast to the WT, which showed a single, short, non-septate germ tube giving rise to a mature and functional appressorium. Similarly, unlike in case of the WT, only very few *dam1*Δ appressoria could penetrate and colonise the rice leaf sheath tissue 40 hours post inoculation (Fig. 6C and D). Next, we examined the virulence of the WT and *dam1*Δ mutant on leaf tissue of an alternative (barley) host. While the WT strain developed typical disease symptoms 5 dpi, the mutant showed only superficial vegetative growth. Thus, our results show that *Magnaporthe* Dam1 is involved in differentiation of the infection structure and is crucial for host invasion.

**Figure 6:**
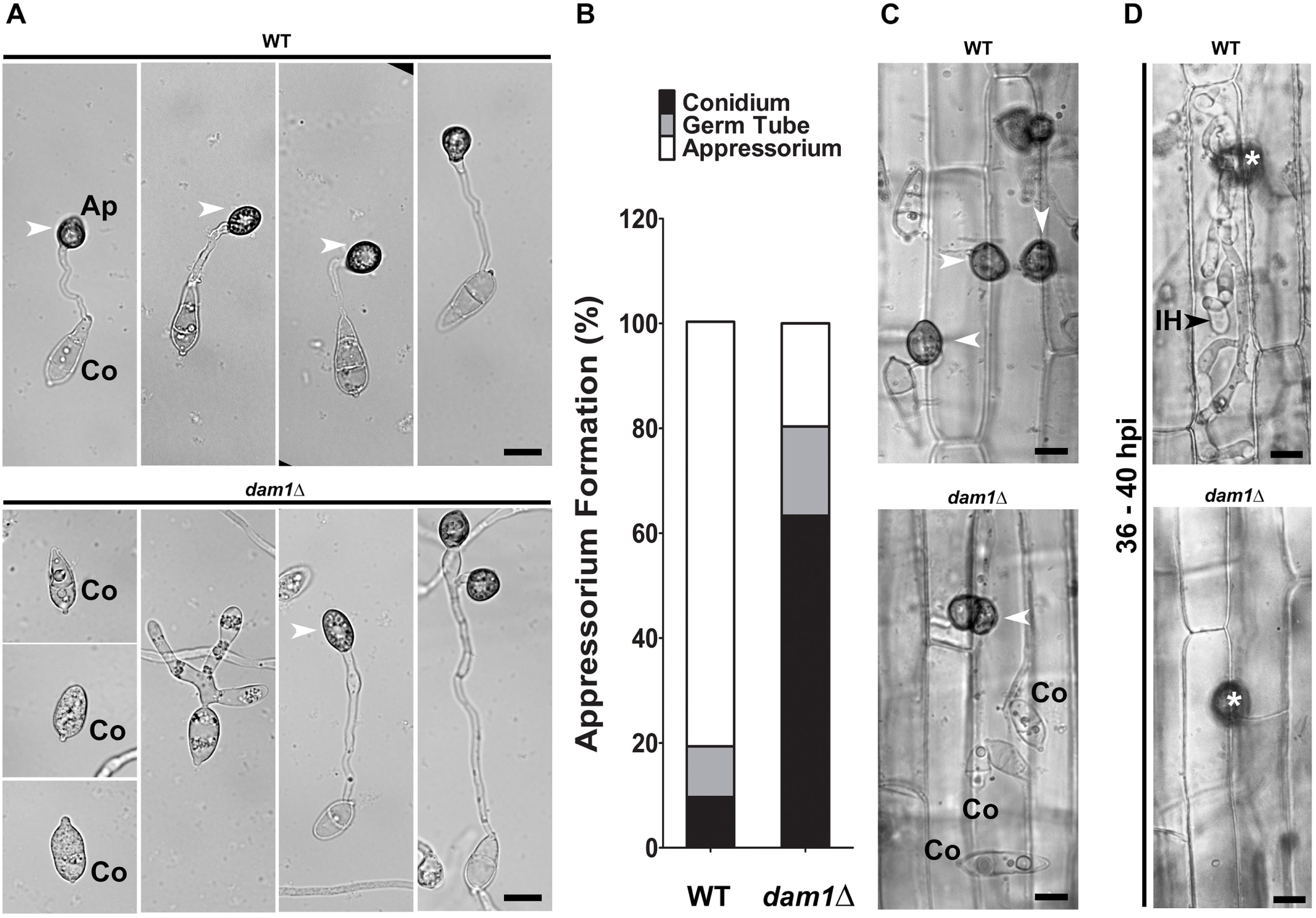
Dam1 function is required for proper pathogenic development. A) Micrographs of pathogenic (appressorium) development on hydrophobic surface (glass coverslips) at 24 hpi. Arrowheads depict appressoria. Co, conidia. Bars = 5 µm. B) Bar chart showing frequency of appressorium formation at 24 hpi on a hydrophobic surface. *P*< 0.05, n = 150 - 200. Data represents mean from three independent experiments. C) Micrographs showing appressorium formation on rice leaf sheaths. Arrowheads depict appressoria. D) Host tissue invasion on rice leaf sheaths inoculated with the WT or *dam1*Δ conidia and observed at 36 - 40 hpi. Arrowheads show invasive hyphae (IH). Asterisks indicate appressorium. Bars = 5 µm.

In summary, our results highlight the importance of dynamics of the outer kinetochore proteins in proper chromosome segregation and polarised growth crucial for the asexual and pathogenic development in *M*. *oryzae*.

## Discussion

A key difference between fungal and metazoan kinetochores is the Dam1/DASH complex that connects the inner KT to the spindle microtubules in fungi. The heterodecameric Dam1 complex is essential for survival in budding yeasts but not in fission yeast. A recent review hints that in the filamentous fungus *Neurospora crassa,* not all Dam1 complex members may be essential (Freitag, 2016), suggesting structural and functional differences in the outer KT in filamentous fungi compared to that of yeasts. In yeasts, whether the Dam1 complex is required for viability is likely dependent on the number of MTs attached to the KT (Burrack et al., 2011, Thakur and Sanyal, 2011). However, the relationship between viability, DASH complex and the number of MTs is not clear in filamentous fungi, where single KT-MT attachment studies are currently lacking. Here, we show that the key members of the DASH complex, Dam1 and Ask1, like the previously studied MoDuo1 (Peng et al., 2011), are not essential in *M*. *oryzae*, suggesting that more than one MT likely binds to a KT in the blast fungus or other interactions stabilise this KT-MT attachment. The fact that we were able to generate deletion mutants of *DAM1* and *ASK1* implies that Dam1 and Ask1 are not essential for viability in *Magnaporthe*. Although there is no complete cell cycle arrest and loss of viability in all cells within the culture, it is likely that only a fraction of cells progress further. This may not be so obvious during multicellular hyphal growth but becomes particularly evident during conidiation and appressorium formation. Furthermore, taking into account the greatly reduced capacity of *dam1*Δ to form conidia and subsequently appressoria that are able to establish infection in hosts, the net capacity for successfully infecting host plants is 2 orders of magnitude lower than that of the wild type. Another feature that differentiates fungal kinetochores from that in humans is the timing and order of assembly of middle and outer kinetochore proteins. In *S*. *cerevisiae* the Dam1 complex is associated with the KT throughout the cell cycle, while all *S*. *pombe* DASH proteins except for Dad1 are recruited to the KT during mitosis. In *M*. *oryzae* we found that, while Dam1 and Ask1 localised to the nucleus at the onset of mitosis and persisted there through chromosome segregation and nuclear migration, the MIND complex protein Mis12 appeared to be a constitutive member of the kinetochore, associating with the nucleus throughout the cell cycle at all stages of fungal development. *S*. *pombe* spindle kinesins Klp5/6 and microtubule plus-end polymerase Mtc1/Alp14 also display metaphase specific kinetochore localisation (Nakaseko et al., 2001, Garcia et al., 2002). Localisation of *M*. *oryzae* Mis12 in the form of a single spot per nucleus before mitosis indicates kinetochore clustering during interphase. These single spots of Mis12 appeared as multiple foci during mitosis, highlighting the dynamic nature of the KT marker protein. These kinetochore dynamics in *M*. *oryzae* are similar to those reported in *S*. *pombe*, where KT de-clustering is seen in metaphase (Goshima et al., 1999). In contrast, de-clustering of KT during mitosis is not visibly evident in the budding yeasts (Roy et al., 2012). Yeast MIND (Mis12) and NDC80 complexes show great plasticity, many copies of proteins being added to the ‘G1 configuration’ of the KT to form the ‘anaphase configuration’ kinetochore (Dhatchinamoorthy et al., 2017). Such a plastic KT structure likely maintains more stable attachment during chromosome separation, while allowing correction of mis-attached chromosomes during metaphase. Whether this is the case in *M*. *oryzae* and can be extended to other outer kinetochore complexes in filamentous fungi will require further study. We observed that the size of the *M*. *oryzae* Mis12 cluster associated with the nucleus before mitosis was larger than the one observed after division (Fig. 1B), likely due to a duplicate set of kinetochores associated with each nucleus or assembly of additional Mis12 complexes prior to mitosis. Overall, the cell cycle dynamics of DASH complex proteins and Mis12 in *M*. *oryzae* are more similar to those in *S*. *pombe* than in budding yeast or humans.

The Dam1 protein in fission yeast is involved in timely anaphase onset and lack of Dam1 leads to lagging chromosomes and sister chromatids occasionally segregating to the same pole (Sanchez-Perez et al., 2005). Along with the spindle kinesin Klp5, Dam1 is involved in chromosome biorientation. In budding yeast, the Dam1 complex proteins play an important role in maintaining spindle structure and integrity, with *dam1* mutants showing diverse spindle defects ranging from elongated, hyperelongated, extremely short to even broken spindles (Hoffmann et al., 1998, Janke et al., 2002, Cheeseman et al., 2001a, b). Similarly, mutations in budding yeast motor proteins, such as Kar3 (Kinesin-14/Klp2), Cin8, Kip1 (kinesin-5) and Kip3 (Kinesin-8/Klp5/Klp6) involved in microtubule stability, also display changes in spindle length (Straight et al., 1998, Zeng et al., 1999). Further, Dam1 plays a role in correct kinetochore attachments and bi-orientation in budding as well as fission yeast, lack of which leads to activation of the spindle assembly checkpoint (SAC), delaying the onset of anaphase (Janke et al., 2002, Sanchez-Perez et al., 2005, Buttrick et al., 2012). Dam1 phosphorylation by Aurora B/Ipl1 kinase in budding yeast and Polo kinase Plo1 in *S*. *pombe* allows dissociation of incorrect KT-MT attachments. This KT is now free to be captured by a new MT until bi-orientiation of sister chromatids is sensed by tension between the two spindle poles. The *M*. *oryzae* Dam1 sequence also shows the Polo consensus site DTSFVD (amino acids 150-155). In *Magnaporthe*, loss of Dam1 function led to delayed anaphase onset, sometimes ending in unequal nuclear division. The delay may be due to SAC activation as a result of lack of chromosome bi-orientation. These events of improper nuclear division and cell cycle arrest account for the reduced hyphal growth and loss of viability of conidia. Further the *dam1*Δ mutant showed altered spindle length with certain instances of collapsed spindles. Thus, *Magnaporthe* Dam1 likely plays a role in both maintenance of spindle structure as well as bi-orientation of chromosome.

In *M*. *oryzae*, differentiation of germ tube into appressorium and conidiophore into conidium are morphologically similar processes requiring a switch-over from polarised to isotropic growth associated with asymmetric cell division. While the role of cell cycle checkpoints in appressorium development has been worked out, similar studies during the development of the aerial conidiophore structure have proved technically more challenging so far in *Magnaporthe* and needs more attention. Two different types of cytokinesis have been described in *M*. *oryzae* depending on the site of septation– 1) septation at the site of mitosis in growing hyphae, and 2) septation spatially uncoupled from mitosis during appressorium formation, where the septum is placed at the neck of the newly formed appressorium while the mitosis occurs in the germ tube (Saunders et al., 2010). We observed both these types of cytokinesis during conidiation. The first septation occurred at the neck of the incipient conidium, away from the site of nuclear division which occurred in the conidiophore stalk, similar to the one seen during appressorium development. The deposition of the two subsequent septa in the developing conidium occurred at the site of mitosis as seen in the the case of vegetative hyphae. We further showed that nuclear association of Dam1 plays a crucial role in these three rounds of mitosis during conidial development. Conidial development was significantly impaired in the *dam1*Δ mutant with altered septation most likely due to aberrant microtubular dynamics and/or unequal nuclear division. Another DASH complex protein in *M*. *oryzae*, MoDuo1, has previously been implicated in proper conidiation; however, its precise role in the process remained elusive (Peng et al., 2011). Defective septation, similar to the one during conidial development in the *dam1*Δ mutant, has also been found in the absence of MoTea4 function, which is mainly involved in cell polarity in *M*. *oryzae* (Patkar et al., 2010), suggesting a non-canonical role for *M*. *oryzae* Dam1 and Ask1 during polarised growth. Indeed, loss of Dam1 function led to defects in hyphal morphology and patterning, with the *dam1*Δ mutant exhibiting excessive branching, irregular hyphal diameter and altered cell size. In *M*. *oryzae* loss of MoSpa2, a component of the spindle pole body, also results in excessive branching (Li et al., 2014). Interestingly, apparently independently of its role as a kinetochore protein, dynamic DASH complex punctae of varying sizes were seen at the hyphal tips under polarised growth during interphase. In *Aspergillus nidulans* hyphal tips, most MTs are arranged with their plus-ends directed towards the tip (Konzack et al., 2005). Further, kinesin KipA moves along microtubules towards the hyphal tip and accumulates at the MT plus-ends, and loss of kipA affects polarity maintenance due to changes in MT-cortex interaction during hyphal growth (Konzack et al., 2005). Taking into account that DASH complex proteins are plus-end MT binding proteins in yeasts, the tip signal in *M*. *oryzae* is likely from Dam1 associated with the MTs at the plus ends, driving hyphal extension. The localisation pattern of Dam1 and Ask1 in *M*. *oryzae* is also similar to the kinesin KipA and microtubule polymerase AlpA in *Aspergillus nidulans*. The Dam1 or Ask1 punctae at the tips were highly dynamic, oscillating between the hyphal end and nucleus, suggesting a possible role in scanning until the nucleus is ready to undergo division. In *S*. *pombe*, Dam1 punctae move along the cytoplasmic MTs (cMTs), occasionally merging into larger oligomers, or crossing over to neighbouring MT tracks (Gao et al., 2010). Further, Dam1 also alters the rate of depolymerisation of spindle as well as cMTs. It is likely that in *M*. *oryzae* Dam1 is associated with the cytoplasmic microtubules during interphase. Indeed, the dynamic Dam1 punctae largely disappeared during interphase upon treatment with the MT-destabilising agent nocodazole. It would be worth studying the *M*. *oryzae* Dam1 interactions with other microtubule-associated proteins and motor proteins and their role in regulating the MT network, especially during interphase.

Penetration peg formation during host invasion involves polarised growth and one round of mitosis in the appressorium, to contribute a nucleus to the emerging invasive hyphae (Jenkinson et al., 2017). Although defects were observed in host penetration and colonisation in the *dam1*Δ mutant, it is not clear whether the defects were only due to aberrant mitotic progression or a combined effect of impaired penetration peg polarity and development. Determining Dam1 localisation patterns during these early infection stages will allow a better understanding of its role in pathogenicity. Here, we propose that in addition to its role in chromosome segregation, Dam1 plays a significant role in polarised filamentous growth. To study whether such a non-canonical function is conserved in other filamentous fungi, further characterisation of respective kinetochores would be required. Since model fungi such as *N*. *crassa* and *Aspergillus*, unlike *Magnaporthe*, are multi-nucleate, detailed studies on DASH complex proteins in these fungi might shed light on some interesting and novel mechanisms underlying the KT dynamics.

Overall, the DASH complex proteins Dam1 and Ask1, though not essential for viability, have important roles to play in cell cycle progression during fungal development, and deletion of *DAM1* drastically reduces the infectivity of the fungus. Lastly, given its importance in pathogenesis and specificity to fungi, Dam1 makes a potential target for a novel antifungal strategy.

## Materials and Methods

### Fungal Strains, Culture and Transformation

*Magnaporthe oryzae* B157 strain (MTCC accession number 12236), belonging to the international race IC9 was previously isolated in our laboratory from infected rice leaves (Kachroo et. al., 1994). The fungus was grown and maintained on Prune Agar (PA). Liquid Complete Medium (CM) was used to grow biomass for DNA isolation, etc. Vegetative growth of the deletion mutants was measured in terms of colony diameter on PA. For conidiation, cultures were grown on PA for 3 days in the dark and then kept under constant light until harvesting. For harvesting conidia, protocol was followed as described (Patkar et al., 2012). Quantification of conidia was done using a haemacytometer. For gene tagging and deletion, plasmids were transformed into *M*. *oryzae* by protoplast transformation or *Agrobacterium tumefaciens* mediated transformation (ATMT) (Mullins et al., 2001). The transformants were selected on YEGA with 300 μg ml^−1^ Zeocin or 300 μg ml^−1^ Hygromycin or Basal medium with 100 μg ml^−1^ Chlorimuron ethyl or 50 μg ml^−1^ Glufosinate Ammonium. The selected transformants were screened by locus specific PCR and microscopy. Single site-specific integration was confirmed by Southern hybridisation.

### Plasmid construction for tagging of genes

Ask1 and Dam1 were tagged with GFP using Marker fusion tagging (Lai et al., 2010). For N-terminal tagging of Dam1, the *DAM1* promoter was amplified using primers Dam1-Pro-F/Dam1-Pro-R from B157 genomic DNA and cloned into p718 at EcoRI/SpeI to obtain p718-Dam1Pro. The 1052 bp *DAM1* ORF and 3’UTR fragment was amplified using primers Dam1-ORF-F/Dam1-3UTR-R and cloned in p718-Dam1Pro in frame with *BAR-GFP* to give p718-GFPDam1. For C-terminal tagging of Ask1, the 3’end of ASK1 ORF was amplified using primers Ask1tagcdsF-ER1/ Ask1tagcdsR-Spe1 from B157 genomic DNA and cloned into p718 at EcoRI/SpeI to obtain p718-Ask1Up. The ASK1 3’UTR was amplified using primers Ask1stop3UTRF-Hpa1/ Ask13UTRR-KpnI and cloned in p718-Ask1Up to give p718-Ask1GFP. For C-terminal tagging of Mis12, the Mis12 3’UTR was amplified using primers Mis123UTRF-H3/ Mis123UTRR-Pvu1 from B157 gDNA and cloned into pFGL347 at HindIII/PstI to obtain pFGL347-Mis123’UTR. The last 1kb of the MIS12 ORF was amplified using primers Mis12orf-ER1/ MIs12cdsR-Sm1. This fragment was then fused with GFP ORF by PCR. The fusion product was cloned in pFGL347-Mis12 3’UTR at EcoRI/KpnI to give pFGL347-Mis12GFP. All primers used in the study are listed in Table S2. All clones were confirmed by RE Digestion. PCR was carried out using XT-5 Polymerase and restriction digestion was done using Fast Digest enzymes (Thermo Scientific). To mark the *M*. *oryzae* nucleus, hH1: mCherry tagged B157 strain was generated using the Sulfonylurea Resistance Reconstitution (SRR) vector pFGL959-H1mCherry, a modified version of the pFGL959 plasmid (Yang and Naqvi, 2011) The plasmid carrying a H1:mCherry expression cassette (ccg1 promoter: hH1: mCherry) was moved into wild type *M*. *oryzae* strain B157 by *Agrobacterium tumefaciens* mediated transformation (ATMT). The tagged strain was confirmed by PCR, fluorescence microscopy and Southern hybridisation. The β-tubulin:sGFP tagging construct was derived from pMF309 (obtained from Michael Freitag). The β-tubulin:sGFP cassette with the ccg1 promoter was digested from pMF309 with *Hpa*I/*Sal*I and ligated to KS-HPT at *Hpa*I/*Xho*I to generate KS-HPT-β-tubulin:sGFP. This KS-HPT-β-tubulin:sGFP vector was moved into the hH1:mCherry tagged B157 strain by protoplast transformation and transformants were selected on hygromycin. The Ask1-GFP, GFP-Dam1 and Mis12-GFP constructs were transformed into the H1-mCherry tagged strain by ATMT. Details of all strains generated in the study are provided in Table S3. All molecular biology procedures were followed as described previously (Sambrook et al., 1989).

### Construction of plasmids for deletion of genes

The *DAM1* deletion cassette was generated by double-joint PCR. The 972 bp upstream and 530 bp downstream flanking regions were amplified from B157 genomic DNA and fused with the 1.24 kb Zeocin Resistance cassette by double-joint PCR. This construct was cloned into AMT based plasmid. For ASK1 deletion, the 1080 bp 5’ and 889 bp 3’ flanking regions of the *ASK1* ORF were cloned upstream and downstream of the hygromycin resistance cassette.

### Microscopy

Brightfield and epifluorescence microscopy were performed on Olympus BX51 (Olympus, Japan) or Nikon Eclipse80i (Nikon, Japan) microscope with the 40x ELWD or 100x/1.40 oil immersion objectives using the appropriate filter set. Sub-cellular localisation was studied by laser scanning microscopy on a LSM 700 inverted confocal microscope (Carl Zeiss Inc., USA). The objectives used were either an EC Plan-Neofluar 40X/1.30 or a Plan-Apochromat 63X/1.40 oil immersion lens. GFP and mCherry were imaged with the 488 nm and 555 nm laser respectively. For live-cell imaging, fungal cultures were inoculated on glass-bottom petridishes. To study protein dynamics, fungal structures were captured as a time series of z-stack images. The images were acquired through the ZEN 2010 software with the Zeiss AxioCam MR Camera and processed and analysed using ImageJ (https://imagej.nih.gov/ij/download.html) and Adobe Photoshop CS6 software. 3 μg ml^−1^ Calcofluor White (Whitener 28, Sigma-Aldrich, USA) was used to stain cell wall and septa of conidia. To examine the effects of the microtubule inhibitor nocodazole on GFP-Dam1 dynamics in germ tubes, germinating conidia were treated with 0.5 μM nocodazole for 15 minutes and images were taken every 5 minutes.

### Pathogenicity Assays

For appressorial assays, conidia were harvested from 10-day-old prune agar cultures. Aliquots (20 μL) of conidial suspensions (5×10^4^ conidia/mL in sterile water with Streptomycin) were applied on hydrophobic cover glass and incubated under humid conditions at room temperature. Conidial germination and appressorium formation were examined 24 h post innoculation. The percentage of appressoria formed was calculated. For penetration assays, rice leaf sheath inoculation assays were performed with conidial suspensions as described (Kankanala et al., 2007) and assessed 36-40 hrs post inoculation. Penetration pegs and infection hyphae were detected by microscopy.

## Acknowledgements

We thank the Bharat Chattoo Genome Research Centre Group for useful discussions and suggestions on the manuscript. We thank Naweed Naqvi (Temasek Lifesciences Laboratory, Singapore) for sharing backbone vectors used to develop gene deletion and tagging constructs in the study. We are grateful to Michael Freitag (Ohio State University, USA) for sharing the pMF309 plasmid used to develop the tubulin-GFP tagging construct.

